# The REFLEX system enables *in vivo* identification of perivascular angiogenic macrophages in the heart

**DOI:** 10.64898/2026.01.27.702189

**Authors:** Tatsuyuki Sato, Takayuki Isagawa, Toshinaru Kawakami, Sosuke Hosokawa, Masamichi Ito, Daigo Sawaki, Shigeru Sato, Yu Nakagama, Kazutoshi Ono, Ariunbold Chuluun-Erdene, Thuc Toan Pham, Ryohei Tanaka, Atsumasa Kurozumi, Hiroaki Semba, Masaki Wake, Shun Minatsuki, Yasutomi Higashikuni, Norio Suzuki, Masataka Asagiri, Hiroshi Harada, Christian Stockmann, Yasushi Hirota, Yasutoshi Kido, Yoshiaki Kubota, Takahide Kohro, Takahiro Kuchimaru, Ichiro Manabe, Issei Komuro, Norihiko Takeda

## Abstract

Direct identification of physically interacting cells *in vivo* remains challenging because conventional interactome analyses infer signaling partners from transcriptomes and cannot reveal which cells are in direct contact. In pressure-overload induced cardiac remodeling, VEGF-A plays a central role in the maintenance of vascular integrity and cardiac function. However, the cell type which produces VEGF-A and how the VEGF-A peptide is delivered to vascular endothelial cells remains unclear. Here, we developed a genetically encoded platform that combines REFLEX mice with HUNTERuni-seq, enabling unbiased detection and transcriptional profiling of the cells that physically interact with vascular endothelial cells. The REFLEX and HUNTERuni-seq approach identified subpopulations of *Vegfa* positive macrophages which we named perivascular angiogenic macrophages (PVAMs). Although the amount of VEGF-A in PVAMs is small, loss of VEGF-A in PVAMs impaired angiogenesis and systolic function during pressure overload. We additionally show that direct contact between PVAMs and endothelial cells is critical for the delivery of VEGF-A to endothelial cells. Conventional interactome analysis predicted that cardiomyocytes as dominant sources of VEGF-A in the heart. However, cardiomyocyte *Vegfa* deletion had no effect on capillary density nor systolic function in a model of heart failure. These results suggest that VEGF-A signaling does not rely on free diffusion through the interstitium and that cellular proximity and physical contact between PVAMs and endothelial cells are the key determinants of effective signal delivery. Together, these findings establish REFLEX and HUNTERuni-seq as a versatile platform for uncovering biologically critical cell-to-cell interactions and provide new insight into intercellular communication in pathological tissue contexts.

## Introduction

Cell–cell interactions underpin tissue development, homeostasis, and disease. However, the direct identification of physically interacting cells within organs remains challenging. Computational interactome analyses, which integrate transcriptomes with ligand–receptor databases, have provided insights into putative signaling partners, but they presuppose unrestricted cytokine diffusion and cannot determine which cells are in direct physical contact^1,2^. Recent advances in spatial transcriptomics partly mitigate this gap by mapping gene expression *in situ*^3,4^. Nevertheless, both imaging-based and sequencing-based spatial transcriptomics are two-dimensional and do not directly report ligand transfer events. Thus, even with spatial maps, inferring which specific neighbors exchange signals remains an open problem.

This limitation is particularly relevant in fibrotic or edematous interstitium, where the extracellular matrix (ECM) and fluid dynamics constrain the mobility of secreted factors. Fibrosis increases collagen deposition and cross-linking, thereby elevating tissue stiffness and reducing interstitial porosity^5,6^. Local tissue edema expands interstitial volume and increases diffusion distance between capillaries and parenchymal cells, further impeding solute transport^7–10^. In parallel, heparan-sulfate proteoglycans (HSPG) within the ECM sequester heparin-binding cytokines, such as vascular endothelial growth factor A (VEGF-A), limiting their effective range^11–13^. Collectively, these features favor contact-facilitated or short-range delivery over long-range diffusion, implying that proximity—not just expression compatibility—often determines whether signaling succeeds in remodeled tissues.

To overcome these barriers, we developed a genetically encoded reporter system, the REFLEX (REcombinase-activated Fluorescent reconstitution by Ligand-mediated cEllular interaction eXtracellulary) mouse, coupled with HUNTERuni-seq (Highlighting Unknown Neighbors Through Extracellular GFP Reconstitution universally and sequencing). This platform enables unbiased identification and transcriptomic profiling of cells that are physically adjacent to a genetically defined population *in vivo*. As a proof of concept, we applied this system to the adult mouse heart, where angiogenic VEGF-A signaling is essential for adaptive remodeling^14–16^. While transcriptome-based interactome predictions emphasized a cardiomyocyte-to-endothelium VEGF-A axis, and although cardiomyocytes produced a vast majority of VEGF-A transcripts within the heart, their deletion had no effect on capillary density nor systolic function. REFLEX/HUNTERuni-seq directly resolved proximate cellular partners and identified perivascular angiogenic macrophages (PVAMs) as the critical source of VEGF-A. Loss of macrophage-derived VEGF-A impaired angiogenesis and systolic performance under pressure overload, and direct contact between macrophages and endothelial cells was essential for conveying VEGF-A signaling. Together, these findings position REFLEX/HUNTERuni-seq as a broadly applicable platform for revealing biologically critical cell–cell interactions.

## Results

### Generation of the REFLEX mouse enables the identification of physically interacting cells

We previously developed *s*GRAPHIC (secretory glycosylphosphatidylinositol-anchored reconstitution-activated proteins to highlight intercellular connections) system, a vector-based method that labels cells in close proximity. In *s*GRAPHIC, index cells are engineered to secrete a soluble C-terminal fragment of GFP (scGFP), whereas other cells display an N-terminal GFP fragment (nGFP) tethered to the plasma membrane via a GPI anchor. When nGFP-bearing cells are adjacent to scGFP-secreting cells, the two fragments reconstitute extracellularly to restore GFP fluorescence, thereby marking adjacent cell populations (Fig. 1A)^17^. We adopted this system to establish the REcombinase-activated Fluorescent reconstitution by Ligand-mediated cEllular interaction extracellularly (REFLEX) mouse. In REFLEX mice, Cre-negative cells express surface nGFP together with mCherry in the nucleus. When Cre is activated, the nGFP and mCherry cassettes are excised, the cells switch to expressing and secreting scGFP together with EBFP in the nucleus, and the secreted scGFP binds nGFP on neighboring Cre-negative cells to reconstitute GFP (Fig. 1B, C). When crossed with *Cdh5*-CreERT2 mice^18^, cells adjacent to endothelial cells are labeled with GFP in the heart and the kidney (Fig. 1D).

**Figure 1.**
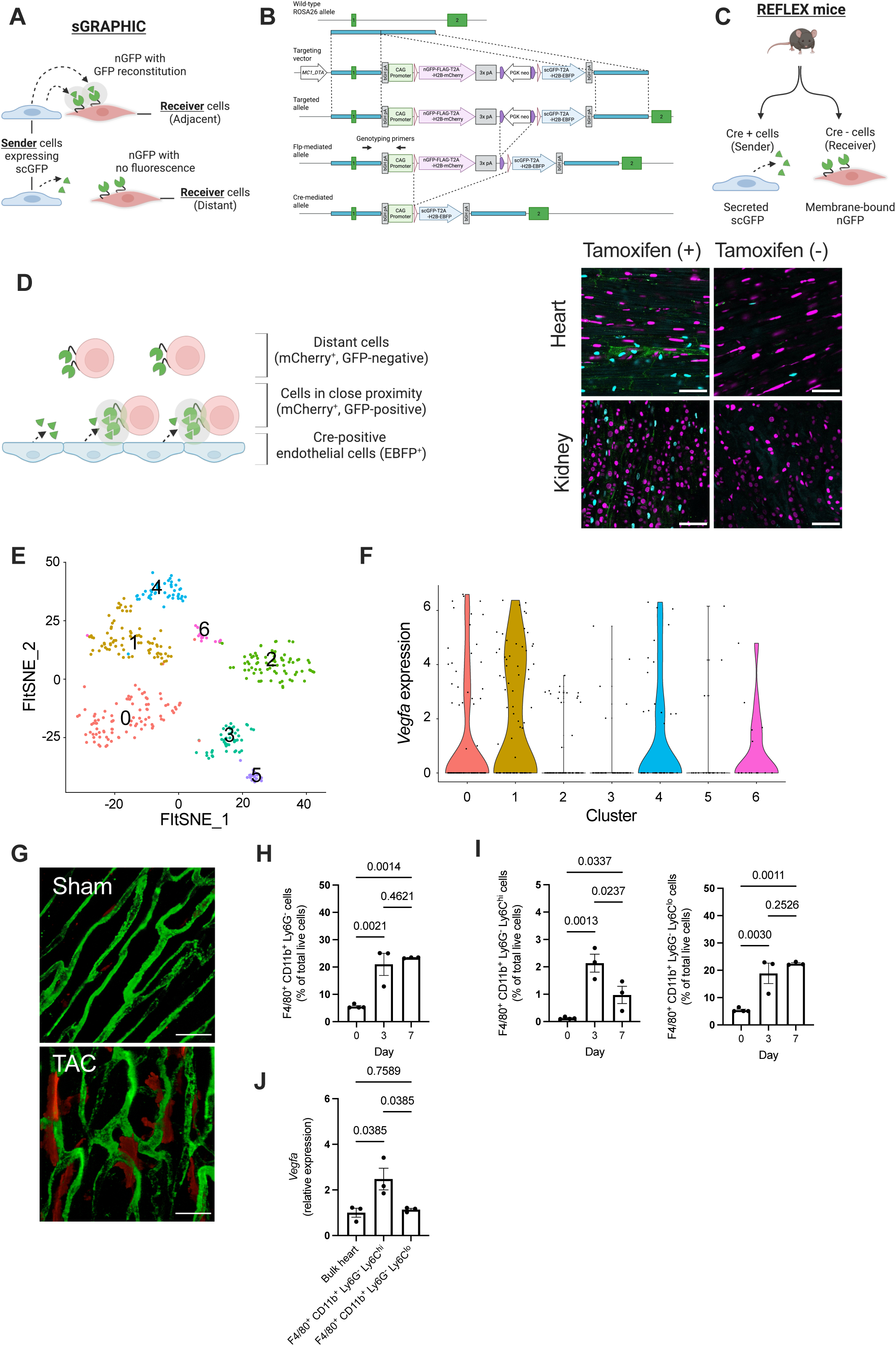
REFLEX mice identify perivascular angiogenic macrophages (PVAMs) as the potential VEGF-A source cells interacting with endothelial cells. A) Scheme of the *s*GRAPHIC system. Sender cells secrete soluble C-terminal fragment of GFP (scGFP), and receiver cells display N-terminal fragment of GFP (nGFP) on their surface. When the receiver cells are in close proximity to the sender cells, GFP is reconstituted and the receiver cells are marked by GFP. Created in BioRender. Takeda, N. (2025) https://BioRender.com/myd7v91. B) Genomic construct of the Recombinase-activated Fluorescent reconstitution by Ligand-mediated cEllular interaction eXtracellulary (REFLEX) mouse. Genotyping primers are shown as arrows. Created in BioRender. Takeda, N. (2025) https://BioRender.com/rn5kfed. C) Scheme of the REFLEX mice. Cells without Cre recombinase express nGFP on their surface together with mCherry. Cre-positive cells express scGFP and EBFP. Created in BioRender. Takeda, N. (2025) https://BioRender.com/9sl26yq. D) Scheme of REFLEX mice crossed with *Cdh5*-CreERTe mice. Following tamoxifen injection, *Cdh5*-expressing endothelial cells express EBFP together with scGFP. Cre-negative non-endothelial cells express mCherry together with nGFP. When the nGFP-expressing cells come in close proximity with Cre-positive endothelial cells, GFP will be reconstituted and the Cre-negative cells will be marked with GFP. The confocal microscopy images show heart and kidney tissues with or without tamoxifen injection in REFLEX *Cdh5*-CreERT2 mice. Scale bar = 50 µm. E) FIt-SNE embedding colored by clusters. Scheme created in BioRender. Takeda, N. (2025) https://BioRender.com/qxomile. F) Normalized expression of *Vegfa* across Clusters presented as a violin plot. G) Confocal microscopy image of the cardiac microvasculature and macrophages 3 days after transverse aortic constriction (TAC). Green: IB4-Alexa488. Red: *LysM*-tdTomato. Scale bar = 20 µm. H-I) The data represent the cardiac infiltration of macrophages (F4/80^+^ CD11b^+^ Ly6G^-^ cells), Ly6C^hi^ macrophages, and Ly6C^lo^ macrophages after transverse aortic constriction (TAC) (n = 3–4 per group). J) Quantitative RT-PCR analysis of *Vegfa* mRNA of bulk heart tissues, Ly6C^hi^ macrophages, and Ly6C^lo^ macrophages collected 3 and 7 days following the TAC procedure (n = 3 per group).

### Perivascular angiogenic macrophages (PVAMs) are identified as endothelial-associated VEGF-A producers by REFLEX/HUNTERuni-seq

We further combined the REFLEX mouse with single-cell transcriptomics, which we named Highlighting Unknown Neighbors Through Extracellular-gfp Reconstitution universally and Sequencing (HUNTERuni-seq). Here, we collect GFP-positive cells from Cre-activated REFLEX mice through cell sorting and perform single-cell RNA sequencing. This allows us to conduct single-cell transcriptomic analysis of cells that reside in functional proximity to the cells of interest.

As a proof of concept, we applied this HUNTERuni-seq to a pathological model of heart failure. We used transverse aortic constriction (TAC). This model induces cardiac hypertrophy accompanied by interstitial fibrosis, edema, and progressive systolic dysfunction^19^. We focused on the vascular endothelium as the index population because endothelial cells are the primary recipients of VEGF-A via VEGFR2 and because microvascular integrity is critical for the maintenance of cardiac function in the failing heart. This setting allows a direct test of whether proximity-based delivery from neighboring cells sustains endothelial support when diffusion of secreted factors is constrained.

Tamoxifen (40 mg/kg body weight) was administered intraperitoneally to REFLEX-*Cdh5*-CreERT2 mice, and the mice underwent TAC three weeks after injection. Three days following TAC, we isolated cells from the heart of REFLEX-*Cdh5*-CreERT2 mice and performed HUNTERuni-seq. We identified seven cell clusters in this HUTNERuni-seq (Fig. 1E), among which Cluster 1 expressed the highest level of *Vegfa* (Fig. 1F). Label-transfer by SingleR annotated 100% of the cells in Cluster 1 as macrophages/monocytes (Supplementary Fig. 1A). Cluster 4, which also showed modest level of *Vegfa* expression, was annotated as monocytes in 97.87% of the cells. Both cluster 1 and 4 on FItSNE showed high expression of macrophage/monocyte marker genes, including *Adgre1*, *Cd68*, *Cx3cr1*, and *Csf1r*. Cluster 1 and 4 expressed low levels of lineage-specific markers for other cell types, including *Cd3d* and *Cd3e* for T cells; *Cd19* and *Cd79a* for B cells; and *S100a8, S100a9,* and *Retnlg* for neutrophils (Supplementary Fig. 1B). Collectively, we have identified clusters 1 and 4 as macrophages/monocytes, both of which express *Vegfa*. Monocytes and macrophages have been reported to be classified into two subtypes, Ly6C^hi^ monocytes/macrophages and Ly6C^lo^ monocytes/macrophages^20,21^. In our data, Cluster 1 more specifically expressed *Ly6c2, Fcgr1*, and *F13a1,* and Cluster 4 expressed more specifically *Itgax*, which is known as a dendritic cell marker, implying that Cluster 1 is Ly6C^hi^ monocytes/macrophages and Cluster 4 is Ly6C^lo^ monocytes/macrophages. Because these Ly6C^hi^ and Ly6C^lo^ monocytes/macrophages are in close proximity to endothelial cells and have significant *Vegfa* expression, we named them type 1 and type 2 perivascular angiogenic macrophages (PVAMs), respectively. The top 10 genes positively correlated with *Vegfa* expression in type 1 PVAMs following exclusion of predicted and uncharacterized genes contain, *Acid sensing ion channel subunit 1*, *Integrin subunit alpha 10*, and *Von Willebrand factor A domain containing 3b*, whereas the list of type 2 PVAMs contain *Human immunodeficiency Virus Type I Enhancer-Binding Protein 1, Mitochondrial carrier 1*, and *Thrombospondin 1*. Type 1 PVAMs were enriched in genes related to tissue remodeling and stress-adaptation, and type 2 PVAMs were enriched in genes related to stress-adaptation, intercellular communication, and angiogenesis. The full list of correlation analysis, including the uncharacterized genes and putative noncoding RNAs, is provided in the supplementary tables 1 and 2.

To validate this finding that PVAMs reside adjacent to endothelial cells, we labeled cardiac macrophages/monocytes using *LysM*-tdTomato mice, which were obtained by crossing *LysM*-Cre mice and LSL-tdTomato mice. We performed TAC on *LysM*-tdTomato mice and found that tdTomato-positive cells reside adjacent to the capillary vessels (Fig. 1G).

Flow cytometry analyses of cardiac tissues following TAC operation showed that cardiac macrophages, defined as F4/80^+^ CD11b^+^ Ly6G^-^ cells, accumulate in the heart after TAC surgery (Fig. 1H). Using the surface marker Ly6C to delineate cardiac macrophage subpopulations^22–24^, we observed that the Ly6C^hi^ subset accumulated predominantly on day 3, whereas the Ly6C^lo^ subset increased on days 3 and 7 (Fig. 1I). While the *Vegfa* gene is highly expressed in Ly6C^hi^ macrophages, the frequency of Ly6C^lo^ cells is higher than Ly6C^hi^, suggesting that both macrophage subpopulations contribute to the production of VEGF-A in the remodeling hearts (Fig. 1J). The representative gating strategy is shown in Supplemental Fig. 2. Collectively, these data suggest that these PVAMs are in direct contact with the endothelial cells and are the potential source of VEGF-A critical for the maintenance of the capillary network and cardiac function.

### PVAMs are required for the maintenance of the capillary bed and cardiac function in the pressure-overload heart

To evaluate the role of PVAMs in cardiac function, we generated myeloid cell-specific VEGF-A knockout mice (mVEGFA CKO: *LysM-Cre*^+/−^ *Vegfa*^flox/flox^) by crossing *LysM*-Cre mice and *Vegfa*^flox/flox^ mice. *Vegfa* expression was significantly suppressed in thioglycolate-elicited peritoneal macrophages (TEPMs) of mVEGFA CKO (Supplemental Fig. 3). We found that the heart weight and steady-state left ventricular ejection fraction were unchanged in mVEGFA CKO mice, indicating that myeloid-cell derived VEGF-A is dispensable in maintaining basal cardiac function (Supplemental Fig. 4). Next, we performed TAC in mVEGFA CKO and control mice and analyzed their hearts after 4 weeks (Fig. 2A). Notably, the whole-heart *Vegfa* levels in mVEGFA CKO were comparable to control mice (Fig. 2B), suggesting that myeloid cell-derived VEGF-A constitutes only a minor fraction of the total VEGF-A in the pressure-overloaded heart. The accumulation of Ly6C^hi^ and Ly6C^lo^ macrophage subsets did not differ between the mVEGFA CKO and control mice (Fig. 2C, D, Supplemental Fig. 5). The left ventricular ejection fraction decreased significantly in mVEGFA CKO (Fig. 2E). Lung congestion was more severe in mVEGFA CKO compared to controls (Fig. 2F). The fibrotic area, heart weight, and cardiomyocyte size were comparable between the knockout and control mice, and the vascular density decreased significantly in mVEGFA CKO mice (Fig. 2G-I). Quantitative RT-PCR analyses revealed no significant changes in the expression levels of *Nppa* and *Nppb,* as well as the ratio of *Myh7 to Myh6,* but a trend towards a decrease in *Atp2a2* expression in mVEGFA CKO mice (Fig. 2J). These results indicate that, although the amount of VEGF-A produced by PVAMs is quantitatively minor, it is physiologically indispensable for adaptive vascular remodeling, contributing to the maintenance of cardiac function in the pressure-overloaded heart.

**Figure 2.**
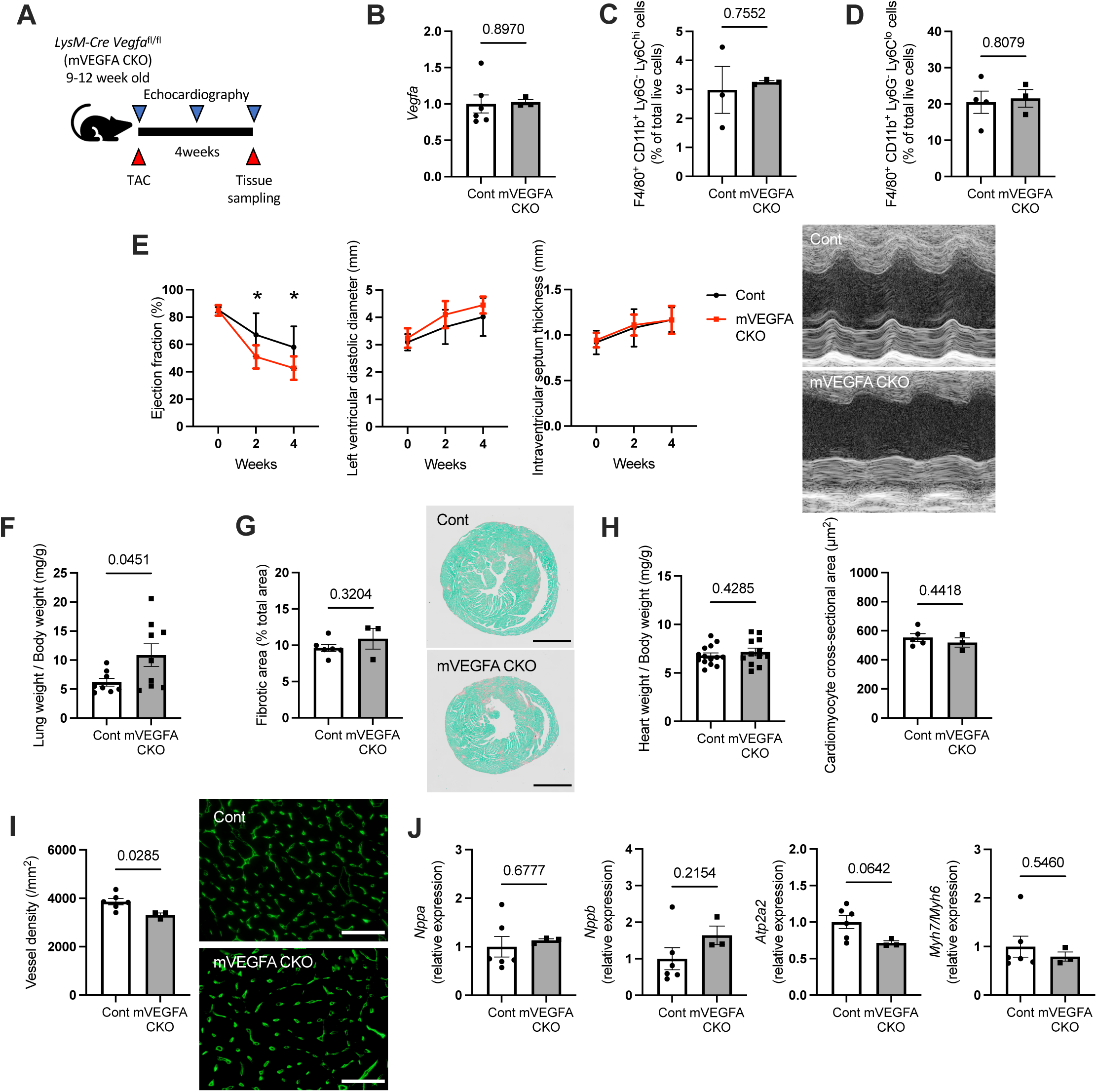
Myeloid cell-derived VEGF-A contributes to maintaining left ventricular systolic function in the pressure-overloaded heart. A) Experimental design. Transverse aortic constriction (TAC) was performed in 12-week-old myeloid cell-specific VEGF-A knockout mice (mVEGFA CKO: *LysM-Cre Vegfa*^flox/flox^; n = 4) and Cre-negative littermate control mice (Cont; n = 3). The heart was harvested 4 weeks after TAC. B) Myocardial *Vegfa* mRNA levels in mVEGFA CKO (n = 3) and control mice (n = 6). C–D) The fraction of Ly6C^hi^ macrophages (F4/80^+^ CD11b^+^ Ly6G^-^ Ly6C^hi^ cells) 3 days after TAC and Ly6C^lo^ macrophages (F4/80^+^ CD11b^+^ Ly6G^-^ Ly6C^lo^ cells) 7 days after TAC in mVEGFA CKO and control mice (n = 3–4 per group). E) Serial echocardiographic evaluation of cardiac function (n = 8–14 per group). * indicates *P* < 0.05. F) Lung weight in mVEGFA CKO and control mice (n = 8–9 per group). G) Fibrotic area of the heart in mVEGFA CKO (n = 3) and control mice (n = 6). Scale bar = 2 mm. H) Heart weight normalized to body weight and myocardial cross-sectional area. (n = 3–14 per group). I) Myocardial vessel density of mVEGFA CKO (n = 3) and control mice (n = 6). Scale bar = 50 µm. J) Quantitative RT-PCR analyses of the mRNA from the hearts of mVEGFA CKO (n = 3) and control mice (n = 6).

### Direct macrophage–endothelial contact facilitates VEGF-A delivery

VEGF-A has three major splice variants: VEGF-A_121_, VEGF-A_165_, and VEGF-A_188_. While the VEGF-A_121_ isoform diffuses freely into the interstitial fluid, VEGF-A_165_ and VEGF-A_188_ possess a heparin-binding domain and adhere to proteoglycans, thus having limited diffusion capacity^25^. We analyzed the abundance of three VEGF-A isoforms in cardiac macrophages and heart tissues three days following TAC. Quantitative RT-PCR using primer pairs for the total VEGF-A and the heparin-binding isoforms showed that the abundance of these isoforms was similar between cardiac macrophages and heart tissues (Supplemental Fig. 6). The results indicate that the composition of each VEGF-A isoform could not account for the cardioprotective effects of myeloid cell-derived VEGF-A.

We next performed an endothelial cell tube formation assay using the mouse endothelial cell line bEnd.3 and the mouse macrophage cell line RAW264.7. A mixed culture of bEnd.3 and RAW264.7 cells in direct contact accelerated tube formation significantly, whereas non-contact co-culture using a cytokine-permeable membrane abrogated the pro-angiogenic effects of RAW264.7 cells (Fig. 3A). Pharmacological inhibition of VEGF-A signaling with sunitinib also suppressed tube formation of bEnd.3 cells (Fig. 3A). These results suggest that direct cell–cell contact between PVAMs and vascular endothelial cells facilitates angiogenic signaling.

**Figure 3.**
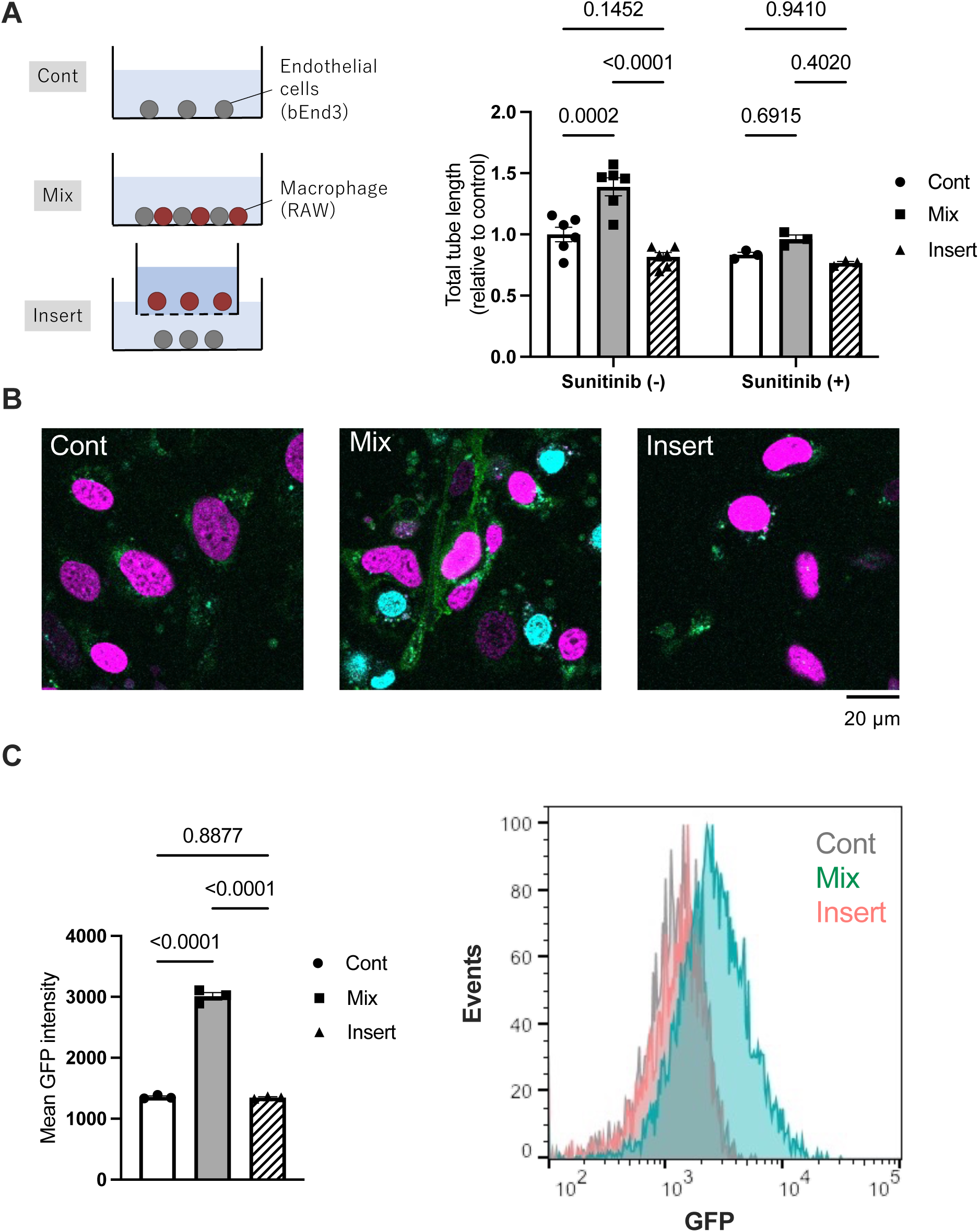
Direct contact between macrophages and vascular endothelial cells accelerates angiogenesis. A) Tube formation by bEnd.3 cells was analyzed with bEND.3 cells alone, with mixed culture with RAW264.7 cells (mix), and with non-contact co-culture (insert) in the presence or absence of sunitinib (1 µM). B) Confocal microscopy images of the cell-cell interaction. bEnd.3 cells cultured alone, mixed with RAW264.7 cells (mix), and non-contact co-culture with RAW264.7 cells (insert). Magenta: bEnd.3 cells. Cyan: RAW264.7 cells. Green: Reconstructed GFP. C) Quantitative flow cytometry analysis of the GFP signal intensity in bEnd.3 cells cultured alone, mixed with RAW264.7 cells (mix), and non-contact co-cultured with RAW264.7 cells (insert). The histogram shows the representative results of each group.

To simulate the delivery processes of PVAM-derived VEGF-A to neighboring endothelial cells, we used the split GFP technique *in vitro*. In this experiment, RAW264.7 cells were transformed to produce soluble GFP fragments and then co-cultured with bEnd.3 cells that express membrane-anchored GFP fragments. When RAW264.7 cell-derived soluble GFP fragments are delivered to the surface of bEnd.3, the GFP signal will be reconstituted in bEnd.3 cells. Confocal fluorescence microscopy and flow cytometry analysis revealed that GFP signals were abundantly detected when RAW264.7 and bEnd.3 cells were co-cultured in the same dish. However, the signal was not detected when they were co-cultured in a non-contact manner (Fig. 3B, C). These data support a model in which PVAMs act as physical carriers delivering VEGF-A to endothelial cells, thereby bypassing diffusional barriers in the remodeled myocardium.

### Conventional interactome analysis fails to detect the source cell of VEGF-A essential for the maintenance of cardiac function

To see whether the conventional interactome analysis can identify an essential cell–cell interaction like our REFLEX/HUNTERuni-seq system, we performed an interactome analysis focusing on VEGF-A-VEGFR2 interaction using a publicly available dataset from the Tabula Muris Senis database^26^ (accession number GSE132042) using CellChat^27^. We found that the cardiomyocyte-endothelial cells axis has the strongest VEGF-A-VEGFR2 interaction, whereas no interaction was found between leukocytes and endothelial cells (Fig. 4A). To examine the functional role of cardiomyocyte-derived VEGF-A, we generated cardiomyocyte-specific VEGF-A inducible conditional knockout mice (cVEGFA CKO) by crossing mice expressing tamoxifen-inducible Cre recombinase specifically in cardiomyocytes (*Myh6-MerCreMer*)^28^ with *Vegfa*^flox/flox^ mice^29^. Deletion of *Vegfa* in cardiomyocytes significantly decreased *Vegfa* mRNA levels in the whole heart, which indicates that cardiomyocytes are the predominant source of VEGF-A in adult hearts (Fig. 4B). Notably, deletion of cardiomyocyte-derived VEGF-A in the adult stage did not affect the basal left ventricular ejection fraction, heart weight, and vessel density of the heart (Supplemental Fig. 7A–C).

**Figure 4.**
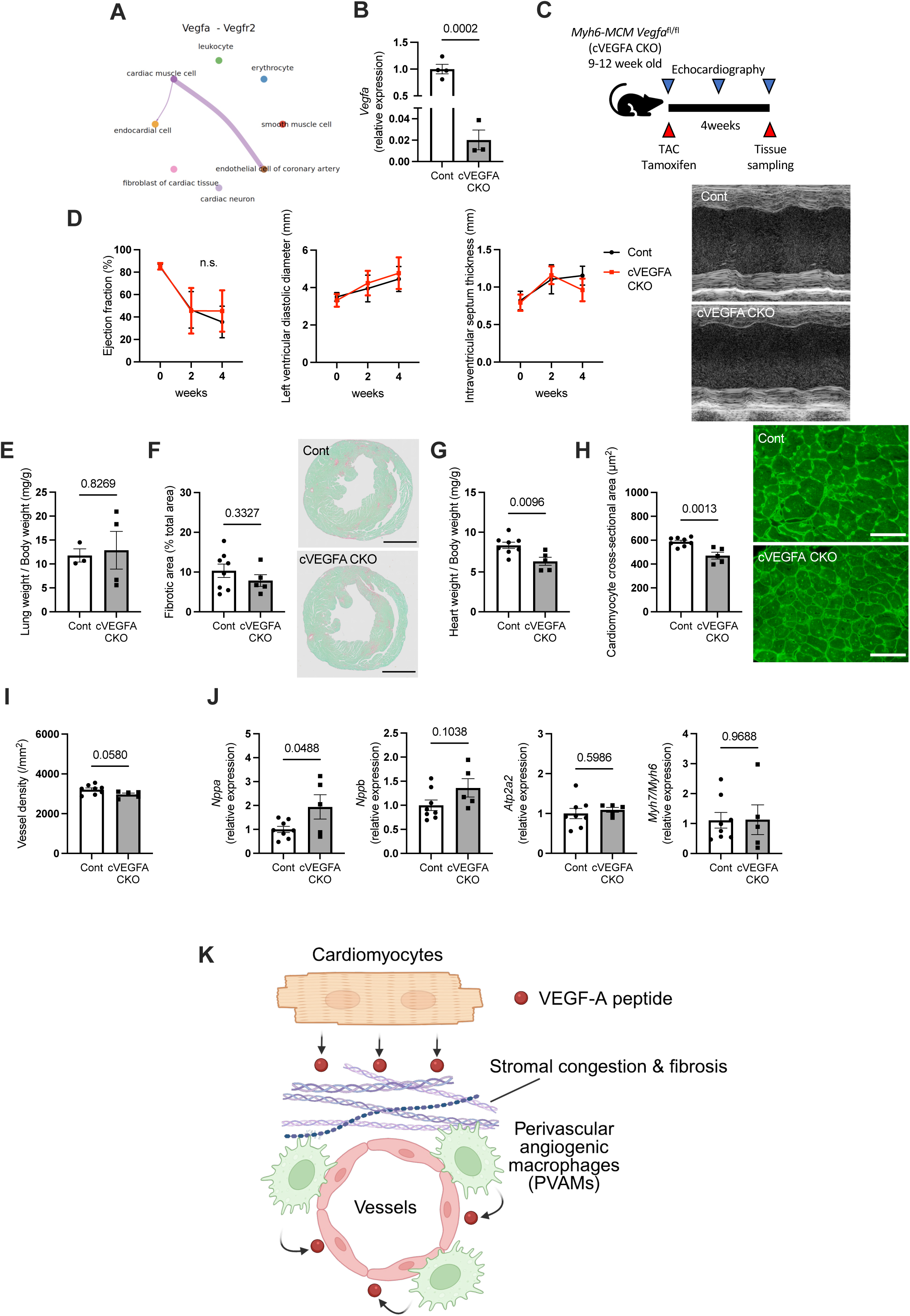
Cardiomyocyte-derived VEGF-A, which was expected to be critical based on conventional interactome analysis, was dispensable for the maintenance of cardiac function and angiogenesis. A) *Vegfa-Vegfr2* interaction in cardiac tissue analyzed using CellChat. B) Quantitative RT-PCR analysis of *Vegfa* mRNA in the heart of cardiomyocyte-specific VEGF-A inducible conditional knockout mice (cVEGFA CKO: *Myh6-MerCreMer Vegfa*^flox/flox^) (n = 3) and Cre-negative littermate control mice (Cont) (n = 4). C) Experimental design. Transverse aortic constriction (TAC) was performed, and tamoxifen was injected simultaneously into cVEGFA CKO and control mice. The heart was harvested and analyzed 4 weeks after the procedures. D) Serial echocardiographic analyses of cardiac function (n = 4–8 per group). E–F) Lung weight and fibrotic area in the hearts of cVEGFA CKO and control mice (n = 3–9 per group) after TAC. Scale bar = 2 mm. G) Heart weight measured in cVEGFA CKO (n = 5) and control mice (n = 8). H) Cardiomyocyte cross-sectional area quantified in cVEGFA CKO (n = 5) and control mice (n = 8). Scale bar = 50 µm. I) Myocardial vessel density in the hearts of cVEGFA CKO (n = 5) and control mice (n = 8). J) Quantitative RT-PCR analyses of mRNA from the hearts of cVEGFA CKO (n = 5) and control mice (n = 8) after TAC. K) Scheme of our hypothesis. Cardiomyocyte-derived VEGF-A fails to reach endothelial cells due to stromal congestion and fibrosis. Perivascular angiogenic macrophages (PVAMs) reside adjacent to endothelial cells and efficiently convey the VEGF-A signal to endothelial cells. Created in BioRender. Takeda, N. (2025) https://BioRender.com/1id3yp1.

Following *Vegfa* deletion and TAC, the left ventricular ejection fraction was similar between cVEGFA CKO mice and control mice (Fig. 4C, D). Lung weight and the extent of tissue fibrosis remained unchanged in cVEGFA CKO mice (Fig. 4E, F). The sizes of the heart and individual cardiomyocytes were smaller in cVEGFA CKO mice compared to control mice, but there was no significant difference in the vessel density between the two groups (Fig. 4G– I), indicating that VEGF-A may act on cardiomyocytes directly and stimulate hypertrophic signal in an autocrine or paracrine manner, but have no effect on angiogenesis^30^. Quantitative reverse transcription polymerase chain reaction (RT-PCR) analyses showed an increase in *natriuretic peptide type A* (*Nppa*) and *natriuretic peptide type B* (*Nppb*) mRNA levels, but no change in the level of *ATPase*, *Ca++ transporting*, *cardiac muscle*, *slow twitch 2* (*Atp2a2)* and *myosin heavy chain7/6* (*Myh7/Myh6*) mRNA ratio (Fig. 4J). Thus, by directly capturing proximity-defined partners in vivo, REFLEX/HUNTERuni-seq identified PVAMs, rather than cardiomyocytes, as the functional VEGF-A donors to the endothelial cells under pressure overload (Fig. 4K). REFLEX mice could be combined with any Cre and inducible Cre lines, providing a scalable strategy applicable for mapping contact-dependent signaling in diffusion-limited tissues.

## Discussion

Conventional interactome analyses infer signaling events from transcript abundance but fail to account for spatial constraints that shape cytokine distribution. We introduce the REFLEX mouse together with HUNTERuni-seq, a platform that directly identifies and profiles physically interacting cells *in vivo* and enables unbiased discovery of functional cell–cell interactions. Using this approach, we discovered that PVAMs, rather than cardiomyocytes, are the critical source of endothelial VEGF-A in the adult heart. Loss of macrophage-derived VEGF-A impaired vascular integrity and cardiac function during pressure overload, despite their small contribution to total *Vegfa* expression.

Notably, cardiomyocyte-specific *Vegfa* knockout ablated the vast majority of cardiac *Vegfa* transcripts, whereas myeloid-specific *Vegfa* knockout left total *Vegfa* transcript levels unchanged. These observations suggest that spatial proximity and physical contact, rather than bulk tissue abundance, are important determinants of effective cytokine delivery in the pressure-overloaded heart. Analogous contact-dependent signaling has been reported in the process of tissue fibrosis, where macrophages adhere to fibroblasts via Cadherin-11 and deliver transforming growth factor-β (TGF-β) directly^31^. The balance between proximity and abundance is likely to vary with tissue architecture, extracellular matrix composition, and receptor availability, and broader application of REFLEX/HUNTERuni-seq will be required to define these rules across organs and cytokines.

These findings, which highlight the importance of PVAM-mediated vascular support and the spatial constraints that govern cytokine delivery, have implications for both anti-VEGF oncology regimens and general heart failure treatment. Anti-VEGF-A antibodies and VEGF-pathway inhibitors are widely used in oncology^32^, and although their use is linked to declines in cardiac function^33^, the pathogenic basis of this toxicity remains incompletely understood. Our data suggest that attenuation of PVAM-derived VEGF-A signaling contributes to this risk, consistent with reports that patients with heightened inflammation face greater odds of heart failure during anti-VEGF treatment^34^. Therefore, source-specific modulation of VEGF-A rather than uniform systemic inhibition may be one solution to mitigating the risk of cardiotoxicity.

In heart failure more generally, therapies that preserve microvascular integrity by maintaining PVAM–endothelial interactions, or that reduce interstitial edema and fibrosis to lessen diffusion barriers, may enhance effective cytokine delivery. By contrast, broad anti-inflammatory or macrophage-depleting approaches could unintentionally compromise vascular support. While cell-type–restricted upregulation of VEGF-A remains challenging, proof-of-concept strategies such as lentiviral transgene expression under macrophage-specific promoters and mannose receptor–targeted delivery have been described^35,36^. Further studies should evaluate targeted augmentation of macrophage-mediated vascular support and delineate patient subsets that would benefit most from such interventions based on inflammatory state and interstitial remodeling.

While the roles of cardiomyocyte-derived VEGF-A in angiogenesis are minimal, cardiomyocytes produce the majority of VEGF-A in adult hearts. Notably, the size of cardiomyocytes in cVEGFA CKO mice was significantly smaller compared with that of control mice. This suggests that cardiomyocyte-derived VEGF-A contributes to cardiac hypertrophy in the adult mouse heart, implying that although cardiomyocyte-derived VEGF-A does not promote angiogenesis, it has an autocrine effect targeting cardiomyocytes themselves. This is somewhat contrary to the previous literature showing that cardiomyocyte-derived VEGF-A is indispensable for angiogenesis and cardiac function in fetal mice hearts^37^. This difference in the function of cardiomyocyte-derived VEGF-A in the fetal/juvenile heart and the adult heart may be explained by the changes in collagen amount and type that occur during aging. A study has shown that the amount of collagen in cardiac tissue increases with age, and a report suggests that the composition of collagen fibers changes during both development and aging^38,39^. VEGF-A contains a heparin-binding domain that interacts with proteoglycans in the extracellular matrix (ECM), and the diffusion capacity of VEGF-A may differ in the adult and fetal/juvenile heart. Previous reports suggest that the composition of ECM dictates cytokine disposition by binding to cytokines and creating immobilized gradients, thereby tuning ligand-receptor interactions, which supports our hypothesis^40^. Soluble VEGF receptor 1 (sFlt-1), which behaves as an endogenous decoy receptor and captures VEGF-A, also has a heparin-binding domain^41^. Therefore, the ECM-sFlt-1 complex may also contribute to the limited diffusion of VEGF-A across the interstitium.

REFLEX/HUNTERuni-seq provides a platform for dissecting direct intercellular communication by genetically defining an index population and fluorescently marking its physically adjacent neighbors for isolation and transcriptomic profiling. The method can be applied to a wide range of cell types, limited primarily by the availability of appropriate Cre drivers, and is well-suited to diffusion-limited microenvironments such as pathologic fibrosis and edema. One limitation of the present study is that the applicability of the system to other organs remains unknown. Although microscopic visualization of other organs from the REFLEX mice, including the bone marrow and the lungs, was not feasible due to high autofluorescence, it is conceivable that HUNTERuni-seq analysis could still be applied to these tissues, as it relies on cell isolation and sequencing rather than direct imaging. Improvement in signal stability and intensity will nevertheless facilitate broader visualization-based applications, which requires further investigation.

In conclusion, cytokine signaling *in vivo* is shaped not only by its abundance but also by the spatial proximity and physical contact between producers and recipients. Using REFLEX/HUNTERuni-seq approach, we have identified a novel cell population, PVAMs, which are critical for angiogenesis and maintenance of cardiac function in the failing heart. Future applications of REFLEX/HUNTERuni-seq include extending analyses to diverse organs and index cell types, quantifying how tissue architecture and ECM remodeling couple proximity to function, and delineating adhesion-dependent interaction networks in both health and disease.

## Methods

### Mice and surgical procedures

Wild-type C57BL/6J mice were purchased from CLEA Japan (Tokyo, Japan). B6.FBV(129)-*A1cf^Tg(Myh6-cre/Esr1*)1Jmk^*/J (JAX stock #005657, *Myh6-*MerCreMer mice), B6.129P2-*Lyz2^tm1(cre)Ifo^*/J (JAX stock #004781, *LysM*-Cre mice), and B6.Cg-Gt(ROSA)26Sor^tm14(CAG-tdTomato)Hze^/J (JAX stock #007914, LSL-tdTomato mice) were purchased from the Jackson Laboratory^28,42,43^. *Myh6-*MerCreMer mice and *LysM*-Cre mice were crossed with *Vegfa*^flox/flox^ mice^29^ to obtain cardiomyocyte-specific VEGF-A inducible conditional knockout mice (cVEGFA CKO: *Myh6-*MerCreMer *Vegfa*^flox/flox^) and myeloid cell-specific VEGF-A knockout mice (mVEGFA CKO: *LysM-*Cre *Vegfa*^flox/flox^). *LysM*-Cre mice were crossed with tdTomato mice to generate myeloid cell-specific tdTomato-expressing mice (*LysM*-tdTomato mice). Mice were housed in a specific pathogen-free facility with a 12-h light/12-h dark cycle. Tamoxifen was used to induce Cre-mediated recombination of the loxP-flanked sites in cVEGFA CKO mice. Tamoxifen was dissolved in corn oil and administered via a single intraperitoneal injection (40 mg/kg body weight)^44^. Male mice aged 7-12 weeks were used for the experiment unless otherwise specified.

The transverse aortic constriction (TAC) operation was performed as follows: First, mice were subcutaneously injected with an anesthetic combination composed of medetomidine hydrochloride (30 µg/mL), midazolam (0.4 mg/mL), and butorphanol tartrate (0.5 mg/mL) dissolved in sterile saline (10 µL/g body weight)^45^. Mice were fixed in the supine position on a heated plate, the chest was opened via a partial sternotomy, and a suture was tied firmly around the 26-gauge needle and the transverse aorta. The needle was then removed, and the chest was closed. Finally, atipamezole, a midazolam antagonist, was subcutaneously injected (1.2 µg/g body weight), and the mice were observed until they were awake. The suture was tied loosely around the aorta for sham surgery, creating no constriction.

### Generation of the REFLEX mouse

To generate REFLEX mice, the CAG promoter and loxP flanked nGFP coding sequence with 3 x poly A signal followed by scGFP coding sequence was inserted into the mouse *Gt(ROSA)26Sor* locus. The generation of REFLEX knock-in mice was performed in TransGenic Inc. (Tokyo, Japan). To construct the targeting vector, a 3.3 kb upstream and a 4.3 kb downstream DNA fragment from the unique *Xba*I site in intron 1 of *Gt(Rosa)26Sor* locus were amplified by PCR from genomic DNA of RENKA ES cell^46^ and used as the homology arms for the targeting vector. These amplified genomic fragments were subcloned into the plasmid vector, which contains the MC1_DTA cassette (polyoma enhancer/herpes simplex virus thymidine kinase promoter driven diphtheriae toxin A gene) as a negative selection marker, with bGH (bovine growth factor) poly A signal to stop ROSA26 endogenous promoter activity, CAG promoter, and bGH poly A signal. To generate the expression unit, the sequence of nGFP-FLAG tag connected with H2B-mCherry by T2A peptide, followed by tandem 3 repeats of SV40 polyA signal (3 x polyA), was flanked by loxP sequences and followed by scGFP connected with H2B-EBFP and bGH poly A signal. A positive selection marker, an FRT-flanked PGK neo cassette (phosphoglycerate kinase I promoter-driven neomycin-resistant gene), was inserted downstream of 3x pA in the reverted orientation. Then, the expression unit was subcloned into the targeting vector downstream of the CAG promoter.

Resulting targeting vector contains MC1_DTA, 3.3 kb 5’ homologous arm, bGH poly A signal, CAG promoter, first loxP site, nGFP-FLAG, T2A, H2B-mCherry, 3 x poly A, frt flanked PGK neo cassette, second loxP site, scGFP T2A H2B-EBFP, bGH polyA, and 4.3 kb 3’ homology arm.

This vector was linearized and introduced into RENKA ES cells (C57BL/6N) by electroporation. After selection using Geneticin, the resistant clones were isolated, and their DNA was screened for homologous recombinant by PCR using the following primer sets: *rosa_F2731*: 5’-CCA TGC TGG AAG GAT TGG AAC TAT GC-3’ and *CAG-R1*: 5’-GAA ACA AGC CGT CAT TAA AC-3’. PCR-positive ES clones were expanded, and isolated DNA from each clone was further analyzed by PCR amplification using the following primer sets: *rosa_F2731* and *CAG-R1* for 5’ homologous recombination and *neo100*: 5’-AGG TGA GAT GAC AGG AGA TC-3’ and *rosa_R10578*: 5’-AAG CTT ACC ATC AAC CTT ATA GTA CAC-3’ for 3’ homologous recombination. Homologous recombination of these clones was also confirmed by genomic Southern hybridization probed with the neomycin-resistant gene sequence.

Homologous recombinant ES cell clones were aggregated with ICR 8-cell embryos to generate chimeric mice. Germline-transmitted F1 heterozygous mice were obtained by crossing chimeric mice with a high contribution of the RENKA background and C57BL/6N mice. The targeted allele was identified by PCR using the primer set *rosa_F2731* and *CAG-R1.* To remove the neomycin-resistant cassette, which may interfere with the expression of the inserted gene, F1 heterozygous mice were crossed with *ROSA26 CAG-Flp* knock-in mice, which express Flp in germ cells. Knock-in allele without the neomycin-resistant cassette was identified with the following PCR primer set, *mCherry-F1*: 5’-ACT ACA CCA TCG TGG AAC AGT ACG AAC-3’ and *splitGFP-R1*: 5’-CAA GAA AGC TGG GTG ACG GTA TCG- 3’. Routine genotyping was performed using the following primer set, *ROSA-F1*: 5’-ACC TTT CTG GGA GTT CTC TG-3’ and *CAG-R5*: 5’-TCG ACC ATG GTA ATA GCG ATG-3’. The generated REFLEX mice were crossed with Tg(CDh5-cre/ERT2) mice (*Cdh5*-CreERT2 mice)^18^ to generate REFLEX-*Cdh5*-CreERT2 mice.

### Confocal tissue microscopy of REFLEX mice

Three weeks after tamoxifen administration in REFLEX mice, the hearts and kidneys were dissected. Whole-mount hearts and sagittal sections of kidneys were then observed using a confocal fluorescence microscope (Leica TCS SP8, Leica Microsystems, Wetzlar, Germany). EBFP was detected with excitation (ex) 405 nm and emission (em) 415-485 nm, GFP with ex 488 nm and em 490-550 nm, mCherry with ex 552 nm and em 600-650 nm. The fluorescence images were analyzed with NIH ImageJ/Fiji open-source software.

### Transthoracic echocardiography

Mice were regularly handled to minimize any stress associated with the echocardiography assessment. Transthoracic echocardiography was performed in non-sedated mice to avoid any cardiosuppressive effects of anesthetic agents. The mice were carefully caught by the examiner’s left hand and placed supine. Data acquisition and analysis were performed every 2 weeks. Images were acquired from a parasternal position at the level of papillary muscles using a Vevo2100® Ultrasound System (FUJIFILM VisualSonics, Toronto, Canada). Left ventricular diameters, ejection fraction, and intraventricular septum thicknesses were serially obtained from grey-scale M-mode acquisition. For each parameter, we averaged values from 3 consecutive cardiac cycles.

RNA isolation, reverse transcription, and quantitative polymerase chain reaction (RT-PCR) Total RNA from the mouse heart was isolated using the NucleoSpin RNA kit (Macherey Nagel, Düren, Germany) according to the manufacturer’s instructions. Complementary DNA was synthesized using ReverTra Ace qPCR RT Master Mix (Toyobo Co., Ltd., Osaka, Japan). Quantitative PCR was performed using THUNDERBIRD SYBR qPCR Mix (Toyobo Co., Ltd.) on a Light Cycler 480 (Roche Diagnostics, Mannheim, Germany). The primers used are described in Supplementary Table 1.

### Flow cytometry and cell sorting

All flow cytometric analyses and sorting were performed using a BD FACSAria Fusion (Becton Dickinson, Franklin Lakes, NJ, USA) and FlowJo software (TreeStar). The isolation of cells from the heart was performed as follows: The heart was cut into small pieces in DMEM containing 10% fetal bovine serum. The minced tissue was then incubated for 2 hours with 1 mg/mL of collagenase type II (Worthington, Freehold, NJ, USA) and 0.74 U/mL of elastase (Worthington). The digested cell suspension was passed through a 23-gauge needle and washed in ice-cold phosphate-buffered saline containing 5% fetal bovine serum. After removing erythrocytes using BD PharmLyse (Becton-Dickinson), blocking with Mouse BD Fc Block (Becton-Dickinson) was performed, and the samples were subjected to fluorescent immunostaining and flow cytometry. The following antibodies were used for the staining: APC-CD11b (M1/70) (eBioscience Inc., San Diego, CA, USA); PE-F4/80 (BM8) (BioLegend, San Diego, CA, USA); FITC-Ly6C (HK1.4) (BioLegend); and Pacific Blue-Ly6G (1A8) (BioLegend).

### HUNTERuni-seq

Tamoxifen (40 mg/kg body weight) was administered intraperitoneally to REFLEX-*Cdh5*-CreERT2 mice, and the mice underwent TAC three weeks after injection. The heart was harvested three days after TAC. Tissue digestion was performed using Cell Dissociation Buffer, enzyme-free, PBS (ThermoFisher, #13151014) instead of collagenase and elastase to prevent the loss of membrane-anchored GFP. After isolation, the cells were biotinylated using EZ-Link Sulfo-NHS-LC-Biotin (Thermofisher, #A39257), followed by barcoding using TotalSeq PE streptavidin A95X and A96X. The cells were then stained using APC/Cy7-CD45 (30-F11) (Biolegend) and BD Pharmingen 7-AAD (BD Bioscience, Franklin Lakes, NJ, USA), and mCherry^+^CD45^+^GFP^+^ cells, mCherry^+^CD45^+^GFP^-^ cells, mCherry^+^CD45^-^GFP^+^ cells, and mCherry^+^CD45^-^GFP^-^ cells were sorted. The cells were then trapped and reverse transcribed using BD Rhapsody (BD Biosciences). The Terminator-assisted solid-phase complementary DNA amplification and sequencing (TAS-Seq) platform was used to perform single-cell RNA-sequencing (scRNA-seq)^47^. Sequencing was performed using the NovaSeq 6000 (Illumina, San Diego, CA) to a depth of 50,000 reads per cell.

### HUNTERuni-seq data processing

Cutadapt-2.1.0^48^ in R 3.6.3 software was used for adapter removal and quality filtering. Gene expression libraries were aligned to mouse Ensemble RNA by Bowtie2-2.4.2, and count matrices were generated using the modified Python script of the BD Rhapsody workflow. Cell barcodes above the inflection threshold of the knee plot of total read counts of each cell barcode identified by the DropletUtils package were identified as valid cell barcodes^49^. Sample origins and doublets were identified based on fold change (FC) of the normalized read counts of the hashtags. The resultant data set was mainly analyzed using Seurat v4.1.3^50^ in R 4.3.1. Doublets were filtered out for quality control. The log-normalized gene counts were calculated using the NormalizeData function (scale.factor = 1,000,000), and highly variable genes were defined by the FindVariableFeatures function (selection.method = mvp, mean.cutoff = c[0.1, Inf], dispersion.cutoff = c[0.5, Inf]). Principal component analysis was performed on the variable genes. Principal components with their P value < 0.05 calculated by the jackstraw method were subjected to cell clustering (resolution = 0.6) and fast-Fourier transform-accelerated interpolation-based t-stochastic neighborhood embedding (FItSNE). Pearson correlation analysis was performed to calculate the correlation of expression between each gene and *Vegfa*. Interactome analysis

The dataset from the Tabula Muris Senis project was obtained from the Cell-x-gene platform and used for the interactome analysis^26^. Initial processing was performed as follows: .h5ad files were converted into a SingleCellExperiment object using the zellkonverter package^51^. The data were filtered based on tissue and developmental stage, and we used the expression data from 3-month-old murine heart tissue. The expression was normalized per 10,000 count and log-transformed. Ensembl ID was transformed into MGI gene symbol using biomaRt. The missing data, null data, and duplicate genes were removed.

Cell-cell communication analysis was performed using CellChat Version 2.2.0^27^. The cells were grouped using the cell annotations included in the dataset. The CellChat object was created from log-normalized expression data and applied to CellChatDB.mouse. Overexpressed genes and cell-cell interaction were identified. Intercellular communication probability was calculated using the triMean method. Finally, we visualized the VEGF pathway and the specific ligand-receptor interaction, together with the extraction of interaction intensity and communication probability.

### Histological analyses of vessel density, cardiomyocyte cross-sectional area, and cardiac tissue fibrosis

The heart was fixed in Tissue-Tek UFIX (Sakura Finetek Japan, Tokyo, Japan) and embedded in paraffin. The embedded tissues were sliced and stained with a combination of biotinylated isolectin-B4 (Vector Laboratories, Burlingame, CA, USA) and streptavidin-FITC conjugate (Vector Laboratories). Alexa Fluor 488 conjugate of wheat germ agglutin (Thermo Fisher Scientific, San Jose, CA, USA) was used for the fluorescent staining of the cell borders to quantify the cardiomyocyte cross-sectional area, while Sirius Red and Fast Green dyes were used for quantifying the degree of tissue fibrosis. The number of vessels and the size of the cardiomyocytes were evaluated in three randomly selected fields per slice and averaged. Bright-field and fluorescent images were acquired using the Virtual Slide System V120 (Olympus, Tokyo, Japan) and FSX-100 (Olympus), respectively. The acquired images were analyzed using NIH ImageJ/Fiji open-source software.

### Whole-mount imaging of the heart

Whole-mount imaging of the cardiac microvasculature and cardiac macrophages was performed as follows. Isolectin B4-FITC conjugate (100 µg/mouse; Vector Lab) was injected via the tail vein to TAC- or sham-operated *LysM*-tdTomato mice. The mice were sacrificed 30 minutes after injection, and the whole heart was visualized using confocal microscopy SP-8 (Leica Microsystems, Wetzlar, Germany).

### Cell culture and *in vitro* tube formation assay

RAW264.7 cells (ATCC, TIB-71) and bEnd.3 cells (ATCC, CRL-2299) were purchased from the American Type Culture Collection (ATCC) (Manassas, VA, USA). Thioglycolate-elicited peritoneal macrophages (TEPMs) were isolated as previously described^49,50^. Thioglycolate was intraperitoneally injected into mice, and the cells were collected from the peritoneal cavity 4 days after injection. The bEnd.3 cell line was stained with 1 mM of Calcein AM solution (Dojindo, Kumamoto, Japan) and was cultured alone, co-cultured with RAW264.7 cells, or with a RAW264.7 cell line seeded on the upper part of a polycarbonate membrane cell culture insert (Merck, Burlington, MA, USA) on a 24-well plate. Before cell plating, the 24-well plate was coated with Geltrex Basement Membrane Matrix (Thermo Fisher Scientific). Sunitinib was purchased from Selleck, Houston, TX, USA. Fluorescent images were obtained using the Operetta CLS High Content Analysis System (PerkinElmer, Waltham, MA, USA) and analyzed using Image J version 2.1.0 software (NIH, Bethesda, MD, USA).

### *SplitGFP* labeling for cell-cell interaction assay

We used *in vitro splitGFP* to capture paracrine interactions^17^. In brief, the lentiviral gene transduction of two types of *splitGFP* reporters, secretory (sc-GFP) and membrane-anchored GFP fragments (n-GFP), was performed into RAW264.7 cells and bEnd.3 cells, respectively. These cells were also labeled with cell nuclei-labeling markers H2B-Azurite and H2B-mCherry, respectively. bEnd.3/n-GFP were cocultured with RAW264.7/sc-GFP either directly or via a polycarbonate membrane cell culture insert with 0.4 μm pores (Corning, NY, USA). The reconstruction of GFP was analyzed using a confocal microscope SP-8 (Leica Microsystems, Wetzlar, Germany) and flow cytometry SH800 (Sony, Tokyo, Japan).

### Statistical analyses

Data are expressed as mean ± standard error. The statistical significance of the differences between the two groups was tested with the Student’s t-test. The Holm-Šidák method was used to adjust the p-value in case of multiple comparisons. One-way Analysis of Variance (ANOVA) or Two-way ANOVA with multiple comparisons using the Holm-Šidák correction was performed to test the difference among more than three groups. All the tests were two-tailed. *P* < 0.05 was considered statistically significant. All analyses were performed with Prism 9 for macOS (GraphPad Software, San Diego, CA, USA).

### Study approval

All animal experiments were approved by the University of Tokyo Ethics Committee for Animal Experiments or the Jichi Medical University Ethics Committee for Animal Experiments. All the procedures strictly adhered to the guidelines for animal experiments of the University of Tokyo or Jichi Medical University.

### Data availability

The scRNA-seq datasets for GFP(+) cells sorted from heart in the REFLEX mouse have been deposited in the DNA Data Bank of Japan BioProject under accession number PRJDB39679. Publicly available scRNA-seq data from the Tabula Muris Senis database (accession number GSE132042) were used for interactome analysis focusing on VEGF-A–VEGFR2 interactions using CellChat.

## Acknowledgments

The authors are grateful to the staff of the Division of Cardiology and Metabolism, Center for Molecular Medicine, Jichi Medical University, for their technical assistance. The authors used ChatGPT (OpenAI, San Francisco, CA, USA) for English language editing and style improvement. All text generated by the AI was carefully checked and revised by the authors to ensure accuracy and originality.

## Author Contributions

T.S., N.T., T.I., T.Ku., and D.S. designed the study, wrote the manuscript, and were involved in all aspects of the experiments, with technical help from Y.N., N.S., M.A., C.S., Y.Ki., and Y.Kub. T.S., K.O., R.T., M.W., H.S., and S.M. were involved in *in vivo* studies; S.S., Y.Hig., and I.M. were involved in the histological studies; S.H., T.Ka., A.K., and M.I. were involved in the bioinformatics analysis; A.C.E., T.T.P., Y.Hir., and I.K. contributed to critical discussion and co-wrote the manuscript.

## Sources of Funding

This study was supported by a Grant-in-Aid for Scientific Research from the Japan Society for the Promotion of Science (JSPS, Japan) (to N.T. 20KK0196, 20K08418, 18H04122, 20H00654, 23K07538, 24KK0157; to T.I. 19K08542; and to T.S. 18J12057, 21J01583, 22K16113; to Y.N. 19K23992; to H.S. 22K08193 and to M.W. 20K17071), and a grant was also received from the Japan Agency for Medical Research and Development (AMED) under grant numbers (JP23gm6210028, JP24gm6510023, JP22jm0110016, JP25gk0210045). N.T. was supported by The Vehicle Racing Commemorative Foundation, The Uehara Memorial Foundation, and The Naito Foundation.

## Disclosures

T.S. has received grant support from The Nakatomi Foundation, The Japanese Heart Failure Society, Fukuda Foundation for Medical Technology, and The Cell Science Research Foundation, and has received honoraria from Sunrise Lab. N.T. has received grant support from Daiichi Sankyo Company. Ltd, Bayer Yakuhin, Ltd, AstraZeneca K.K, Ono Pharmaceutical Co. Ltd, and Bristol Myers Squibb. I.K. has received grant support from Daiichi Sankyo Company. Ltd, Idorsia Pharmaceuticals Japan Ltd, Takeda Pharmaceutical Company Limited, Mitsubishi Tanabe Pharma Corporation, and TEIJIN PHARMA LIMITED., and has received honoraria from ONO PHARMACEUTICAL CO. LTD., AstraZeneca K.K., and Nippon Boehringer Ingelheim Co., Ltd. Y.N. and Y.Ki. have received joint research funding from ROHTO Pharmaceutical Co., Ltd. The other authors declare that they have no competing interests.

---

**Supplemental Figure 1.** A) Annotation of the cells using the SingleR package. B) Mapping of core marker genes onto the FItSNE plot.

**Supplemental Figure 2.** Representative gating strategy for analyzing cardiac infiltration of macrophages (F4/80^+^, CD11b^+^, Ly6G^-^ cells). Cardiac macrophages were divided into two populations depending on the expression level of Ly6C (Ly6C^hi^ and Ly6C^lo^).

**Supplemental Figure 3.** Total RNA was isolated from TEPMs of mVEGFA CKO and control mice. The deletion efficiency of the *Vegfa* gene was calculated using primers that detect the targeted exon (*Vegfa*) and primers for the undeleted exon (*Vegfa*_ex8) for normalization. Data show the mean and the standard error (error bars) of technical triplicates from a representative experiment. N.D., not detected.

**Supplemental Figure 4.** A–B) Echocardiographic analyses and heart weight of myeloid cell-specific VEGF-A knockout mice (mVEGFA CKO) and control mice (Cont) at baseline (n = 3–14 per group).

**Supplemental Figure 5.** A) The fraction of macrophages (F4/80^+^ CD11b^+^ Ly6G^-^ cells) and Ly6C^lo^ macrophages (F4/80^+^ CD11b^+^ Ly6G^-^ Ly6C^lo^ cells) 3 days after TAC (n = 3 per group). B) The fraction of macrophages (F4/80^+^ CD11b^+^ Ly6G^-^ cells) and Ly6C^hi^ macrophages (F4/80^+^ CD11b^+^ Ly6G^-^, Ly6C^hi^ cells) 7 days after TAC (n = 3–4 per group).

**Supplemental Figure 6.** A) Scheme of the primer design. Primer *Vegfa* was designed between exons 2 and 3, which are common in the three major isoforms. Primer *Vegfa*_ex7/8 was designed to detect VEGF-A_165_ and VEGF-A_188_. B) The ratio of heparin-binding domain-containing isoform to total VEGF-A isoforms in cardiac macrophages (n = 3 per group). The results were normalized to the levels of bulk heart tissues.

**Supplemental Figure 7.** A) Tamoxifen was administered at 9 weeks old, and serial echocardiography was performed in cardiomyocyte-specific VEGF-A inducible conditional knockout mice (cVEGFA CKO: *Myh6-MerCreMer Vegfa*^flox/flox^) and Cre-negative littermate control mice (Cont) (n = 3–4 per group). B–C) Heart weight and myocardial vessel density of cVEGFA CKO mice and control mice 4 weeks after tamoxifen treatment (n = 3–4 per group).

**Table 1.**
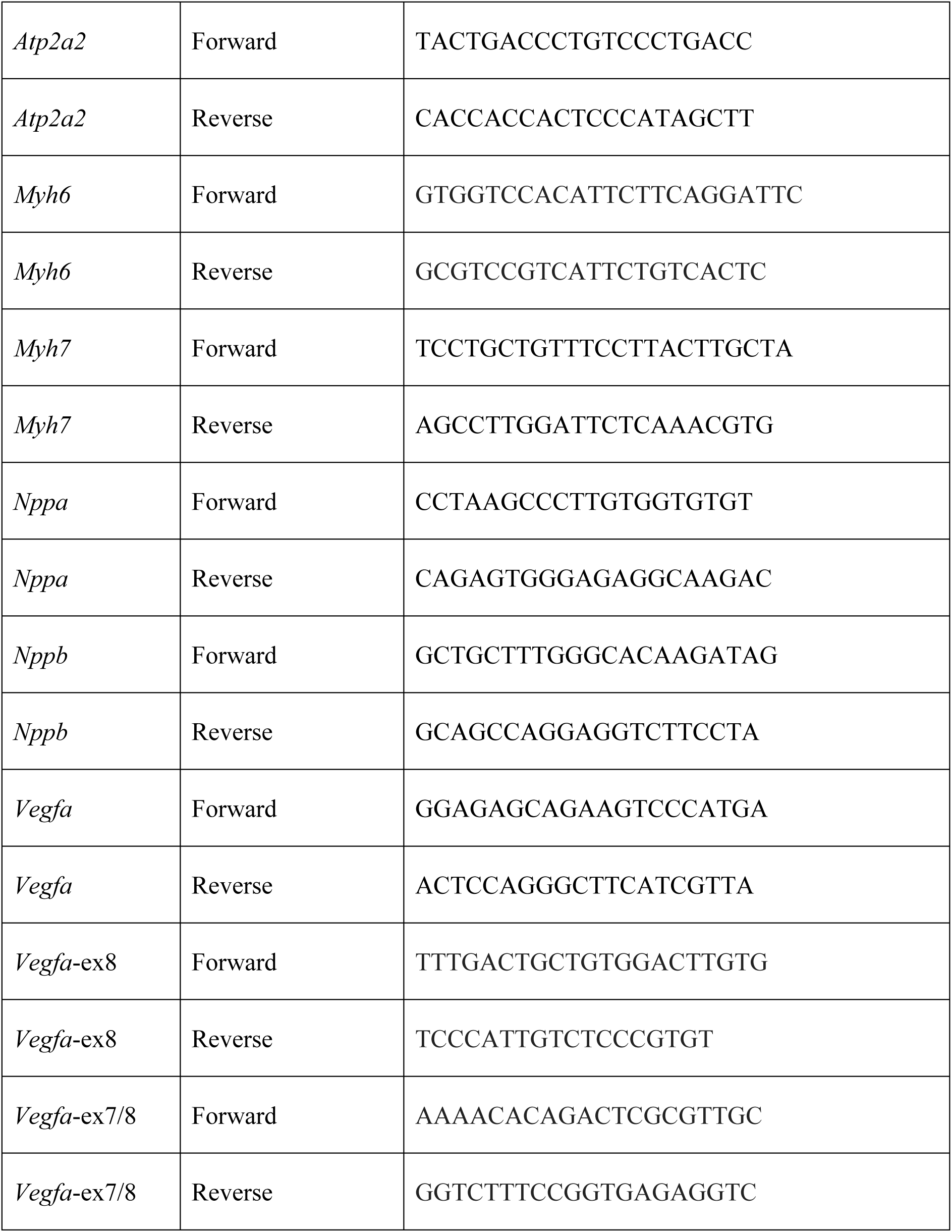
Primers used for the quantitative PCR analyses.

**Table 2.** Top 10 genes the expression of which positively correlated with *Vegfa* in type 1 PVAMs.

| gene | correlation |
| --- | --- |
| *Asic1* | 0.47448515 |
| *Itga10* | 0.39715176 |
| *Vwa3b* | 0.38245409 |
| *Dnajc30* | 0.38150973 |
| *Ppm1g* | 0.37649858 |
| *Acvr2b* | 0.36886327 |
| *T2* | 0.36399639 |
| *Trim46* | 0.36329287 |
| *Abca12* | 0.35511042 |
| *Kctd18* | 0.35363243 |

**Table 3.** Top 10 genes the expression of which positively correlated with *Vegfa* in type 2 PVAMs.

| gene | correlation |
| --- | --- |
| *Hivep1* | 0.65799444 |
| *Mtch1* | 0.64576771 |
| *Thbs1* | 0.63123964 |
| *Dynlt3* | 0.58417796 |
| *Svopl* | 0.57781575 |
| *Fcrla* | 0.55947339 |
| *Upf2* | 0.5565062 |
| *Slc4a4* | 0.55264929 |
| *Gja1* | 0.54838444 |
| *Insrr* | 0.54698871 |

## Notes

https://ddbj.nig.ac.jp/resource/bioproject/PRJDB39679

https://www.ncbi.nlm.nih.gov/geo/query/acc.cgi?acc=GSE132042

