## Supplementary Figures for "The REFLEX system enables *in vivo* identification of perivascular angiogenic macrophages in the heart"

### Supplemental Fig. 1

A

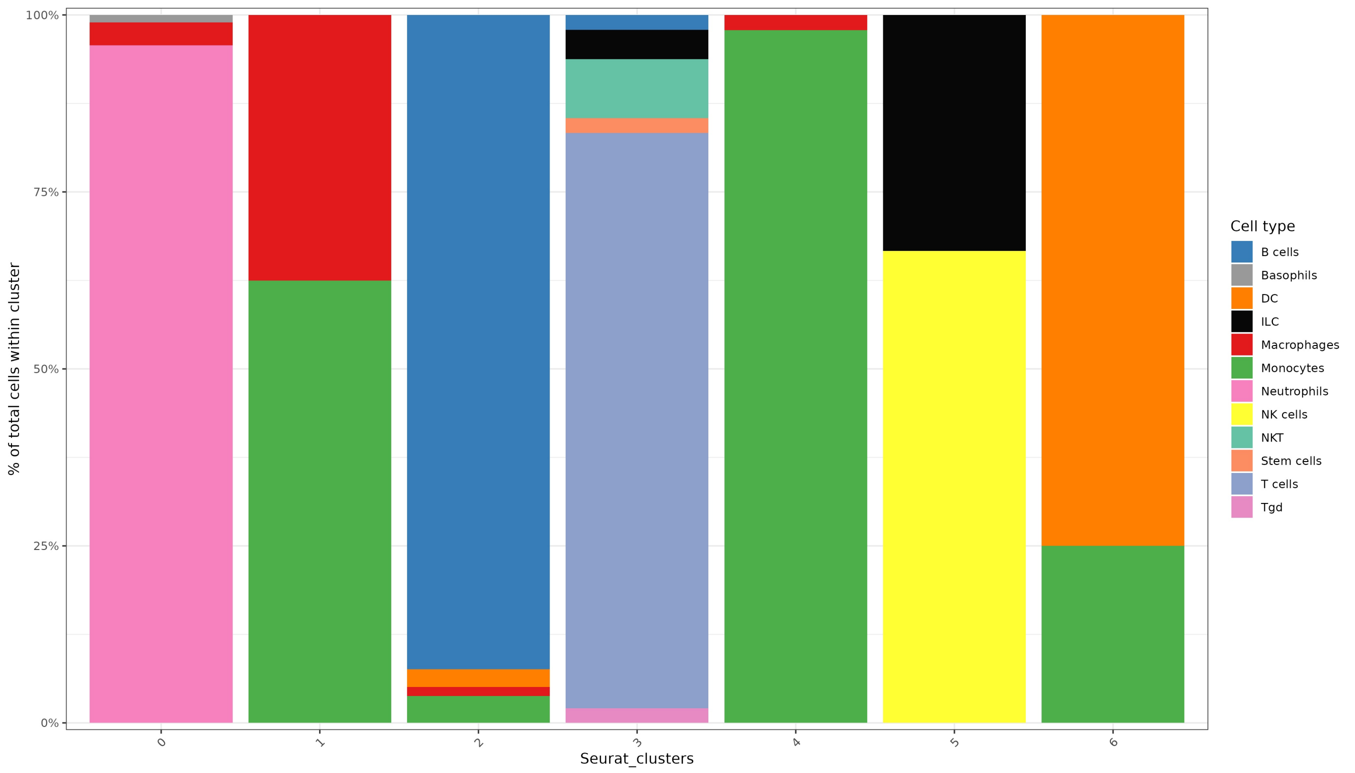


B

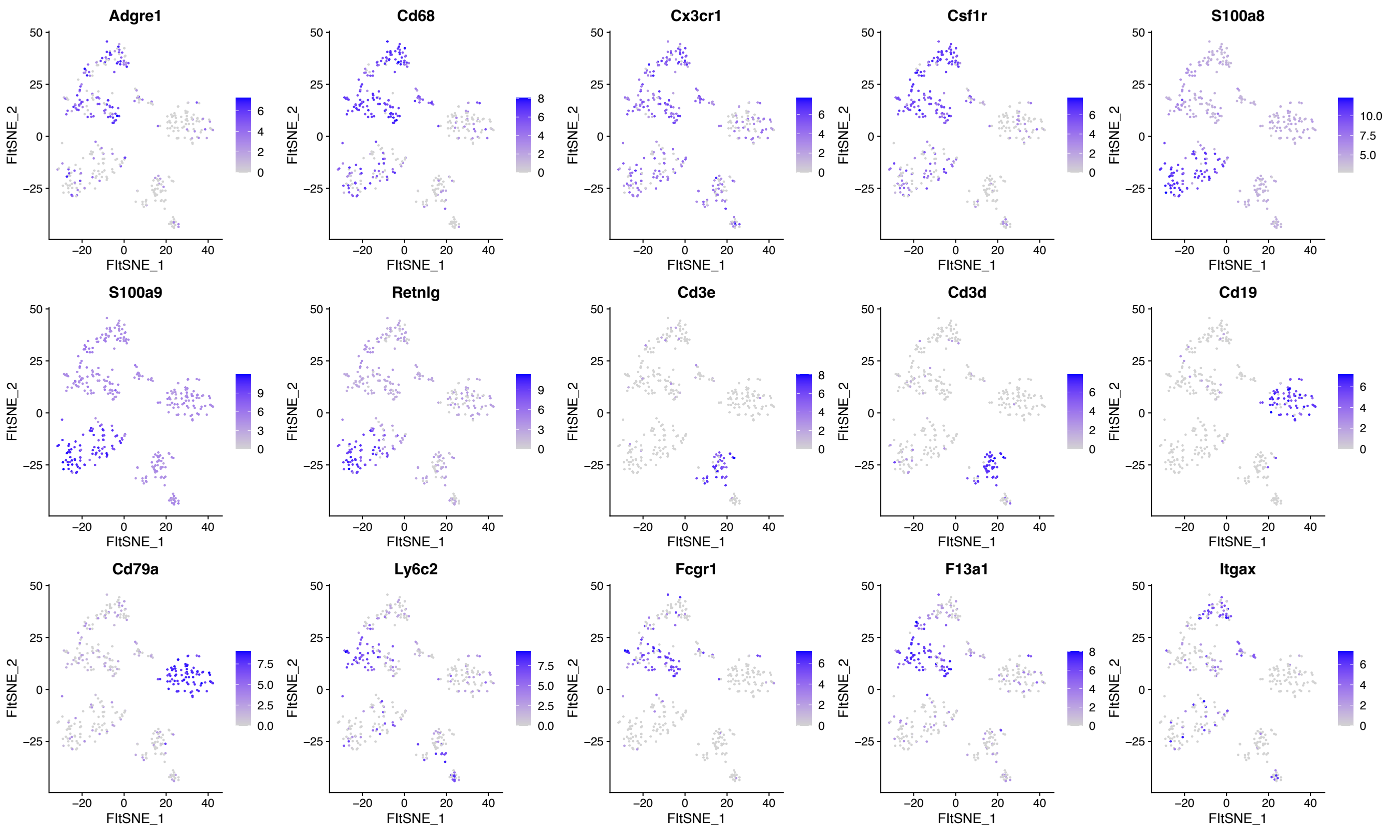

### Supplemental Fig. 2

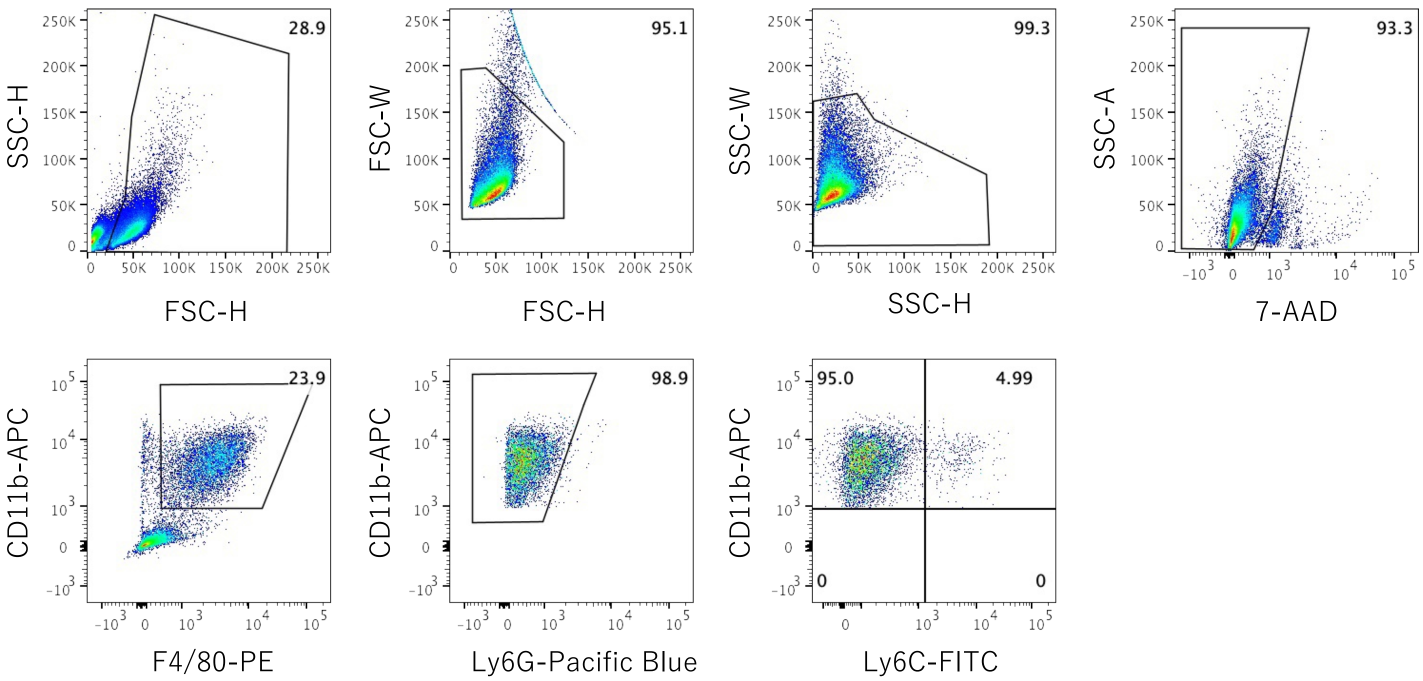

#### Supplemental Fig. 3

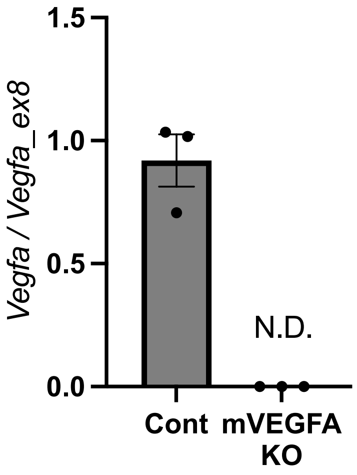

### Supplemental Fig. 4

**A**

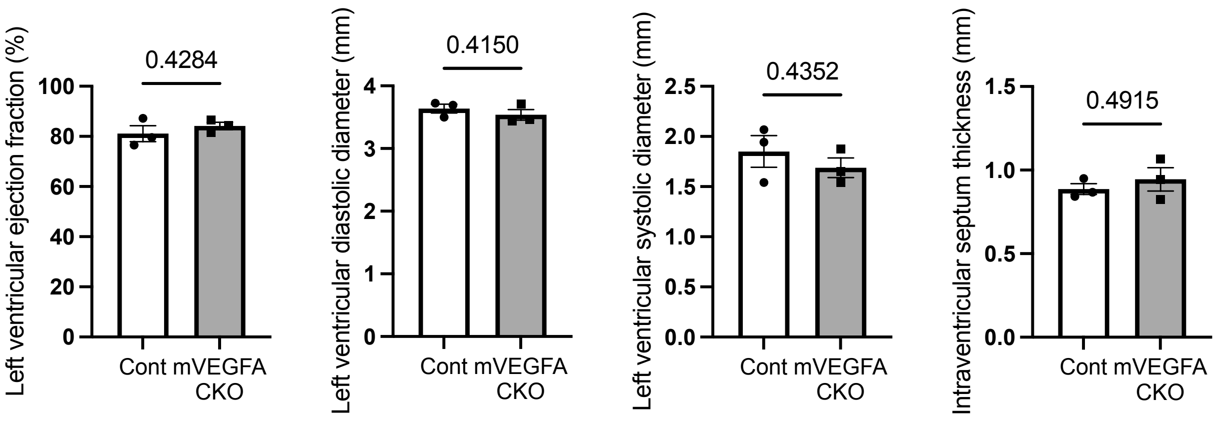


**B**

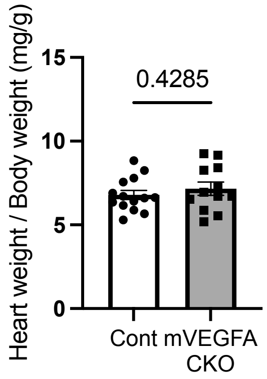

### Supplemental Fig. 5

**A**

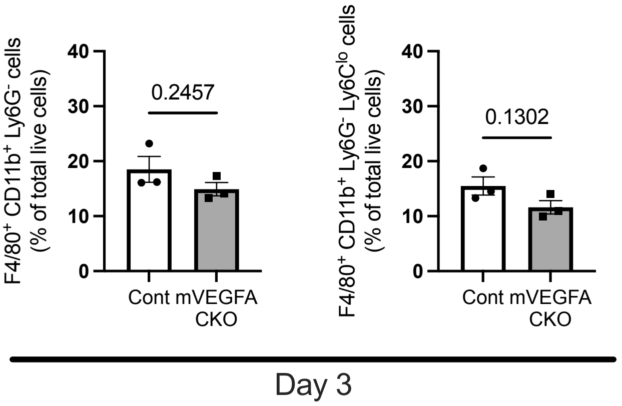


**B**

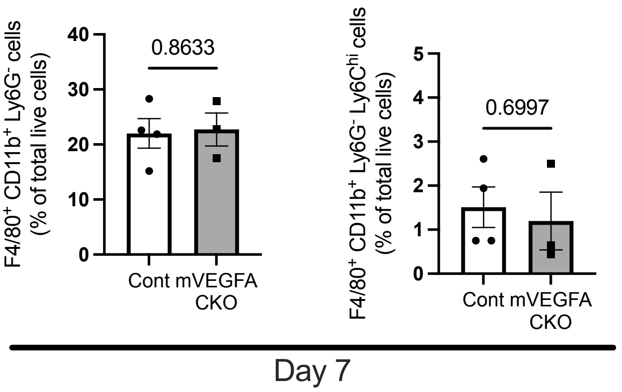

#### Supplmental Fig. 6

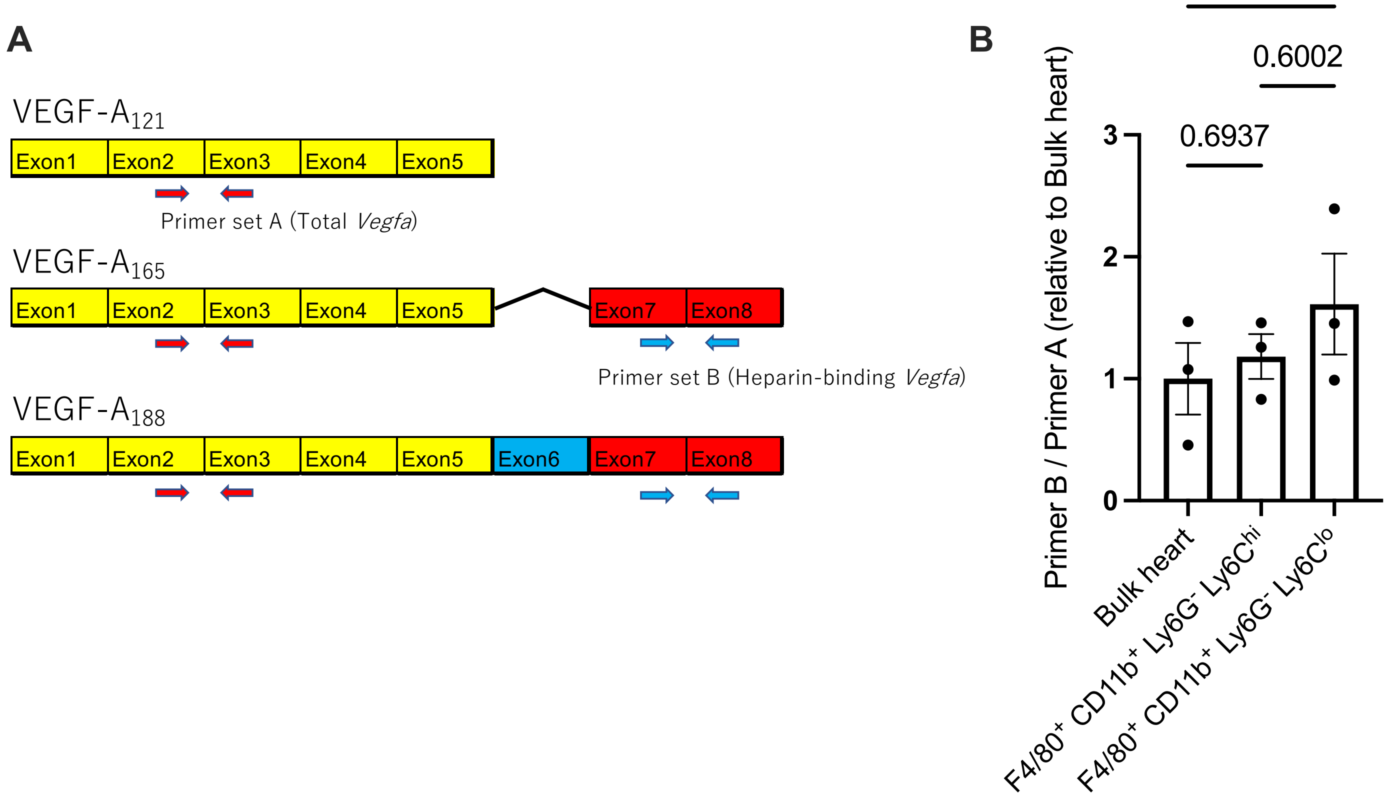

### Supplemental Fig. 7

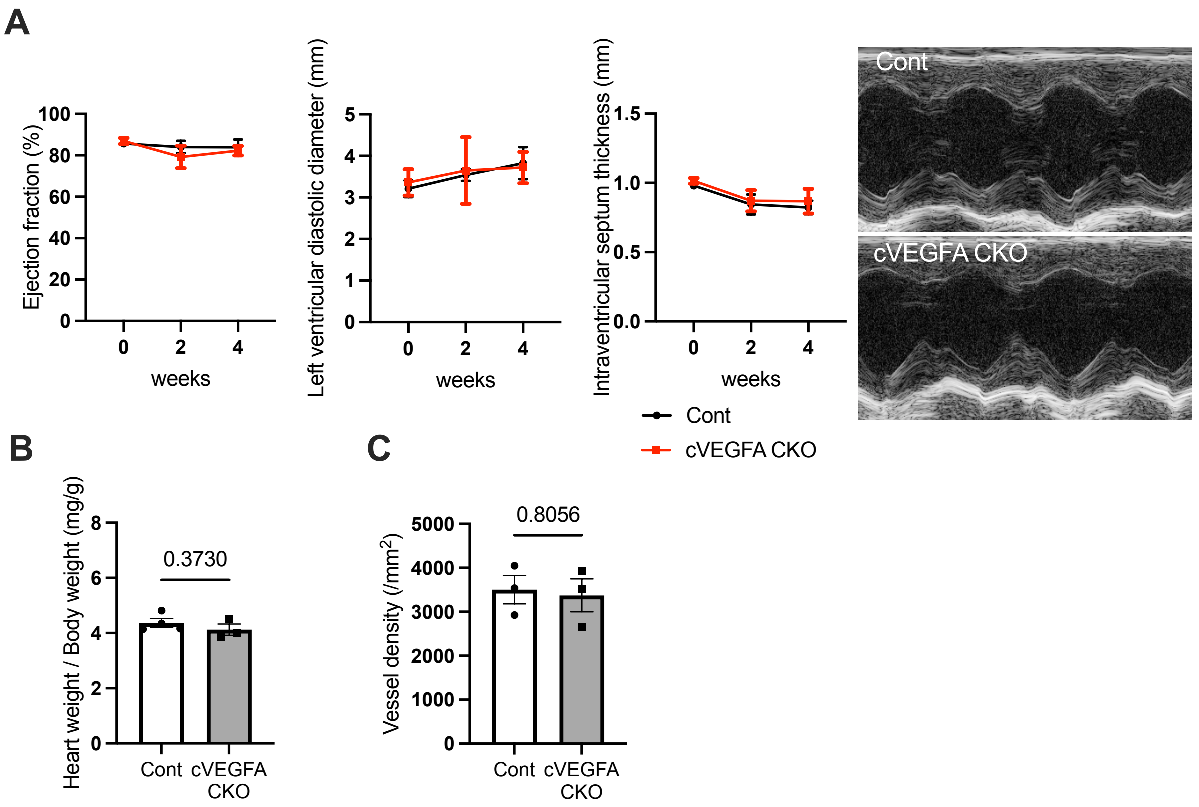
